# Assembly of plant holobionts is governed by nematode communities and their associated microbiota, conditioned by preceding plants

**DOI:** 10.64898/2026.07.02.736003

**Authors:** Holger Heuer, Dirk Schmalowski, Owuraku Amponsah Abu, Marleen Hoernlein, Ute Zimmerling, Jan Reinecke, Katja R. Richert-Pöggeler, Doreen Babin

**Author notes:** Corresponding author: Doreen Babin. University of Göttingen, Institute for Plant Pathology and Plant Protection, Göttingen, Germany (Owuraku Amponsah Abu).

## Abstract

Plants form holobionts by associating with diverse microbiota. Self-organization gives rise to emergent properties of the holobiont, such as increased resistance to pathogens. However, the local factors contributing to the self-organization are not well understood. We hypothesized that nematode communities and their associated microbiota govern the rhizobiome of the model plant holobiont tomato in terms of its suppression of root invasion by the parasite *Meloidogyne hapla*, and that the soil legacy influences the suppressive potential mediated by these biota. In pot experiments, a resistant tomato holobiont was favored by assembly in the presence of a nematode community conditioned by tomato plants, compared to oilseed rape or fallow soil. Nematode communities conditioned by tagetes could enhance resistance even better than tomato. Microbiota from crushed tomato-conditioned nematode communities increased resistance of the tomato holobiont, compared to microbiota of nematode communities conditioned by maize, or heat-inactivated microbiota. The 0.2 µm-filtered microbiota from crushed nematodes had the same effect, suggesting a role of nematode-associated bacteriophages in holobiont assembly. The results indicate that soil nematodes and their associated microbiota play a role in the local organization and stabilization of plant holobionts. They can influence the resistance of plants that subsequently grow in the same soil. From an applied perspective, crop rotation schemes that alter nematode–microbiota communities could be harnessed to engineer crop holobionts.

**SUMMARY STATEMENT:** Depending on plant-specific soil legacies, nematode communities and their associated phages govern holobiont assembly leading to increased resistance against pathogen invasion.

## INTRODUCTION

Harnessing the interactions between plants and their associated biota may be the key for sustainable crop production (Compant et al. 2025). The rhizosphere is the zone directly adjacent to the roots that is strongly influenced by the presence of the plant (Philippot et al. 2013). A specific biotic community, the rhizobiome, comprising bacteria, fungi, protists, nematodes, microarthropods and others, establishes there (Wilschut and Geisen 2021). Plants have coevolved with diverse microbiota, forming discrete ecological units, composed of a host plant, its rhizobiome and other microbial associates, that can be regarded as holobionts (Spooren et al. 2024; Baedke et al. 2020; Mesny et al. 2023). The plant holobiont constitutes a functional network that self-organizes based on interactions among local microbiota, which are connected in complex networks of positive and negative interactions (Mesny et al. 2023). These networks assemble in response to and interaction with the plant. Thereby the plant holobiont assembles through local self-organization, resulting from interactions among a subset of species in the community (including the plant). The biotic interactions constitute a functional network that contributes to the fitness of the members of the holobiont (Bordenstein and Theis 2015). The importance of biotic interactions to stabilize such networks was recently shown for synthetic bacterial communities, where the proteome of the individual members were more determined by the community context than by carbon source, thereby promoting metabolic complementarity and stabilizing the network structure (Moraïs et al. 2026). Due to the high functional redundancy of soil microbiota (Chen et al. 2022), many different beneficial functional networks with varying taxonomic compositions can assemble with the plant to form a holobiont, depending on the soil legacy, the local biotic and abiotic environment, the order of species arrival and other stochastic events. Alternative stable host-microbiota assemblies are well documented for human holobionts (van de Guchte et al. 2018), and should be even more important for the less mobile plant holobionts that assemble from local soil biota. Plants create a soil-borne inheritance of microbiota that have prospered on the holobiont (Spooren et al. 2024). This can benefit the assembly of subsequent plant holobionts growing in the same soil. Self-organization and emergent properties of the holobiont result from local interactions between the organisms involved. An often observed emergent property is an enhanced resistance of the holobiont against pathogens and pests (Spooren et al. 2024). However, the factors contributing to local self-organization of plant holobionts, favoring emergent properties and stability of the system, are not well understood.

Plant-parasitic nematodes pose a global threat to crop production (Savary et al. 2019). Of these, root-knot nematodes (*Meloidogyne* spp.) are considered the most devastating (Jones et al. 2013). However, plant-parasitic nematodes coexist with plants in natural soil up to a certain population density without causing damage (Seinhorst 1965). The juveniles (J2) of these nematodes hatch from eggs in the soil and move towards the roots of a host plant. On the way, they are exposed to the rhizobiome before invading the root. Depending on the presence or composition of the rhizobiome, the reproductive success of plant-parasitic nematodes can be significantly altered (Adam et al. 2014; La et al. 2024). Microbes in the rhizosphere can directly affect the parasites, or indirectly by plant-mediated mechanisms (Topalović et al. 2020b).

Nematodes represent a highly abundant and functionally diverse group of the rhizobiome (van den Hoogen et al. 2019). The structure of soil nematode communities has been associated with soil health and used as an ecological indicator (Gao et al. 2020; Neher and Mosdossy 2025). The plant and its genotype influences the structure of the nematode community (Wang et al. 2023). In turn, the nematode community affects the composition of the microbiome in the rhizosphere and plant performance (Wilschut and Geisen 2021). This suggests that nematodes play a role as part of the plant holobiont. However, the contribution of nematodes to the assembly of the plant holobiont in order to benefit from emergent properties is not well understood.

The cuticle of nematodes is associated with specific bacteria and fungi, depending on the nematode species and soil environment (Elhady et al. 2017; Topalović et al. 2022; Topalović et al. 2019). Diverse genomic sequences of viruses were found in association with nematodes (Huang et al. 2025; Vieira et al. 2022). Recently, it was shown that bacterivorous nematodes can vector phages through soil (van Sluijs et al. 2025). Extracellular phages are highly abundant and diverse in soil (Cobián Güemes et al. 2016). In addition, they have a narrow host range, and thus are likely affecting the bacterial community structure in terrestrial ecosystems (Osburn et al. 2024). It has not been studied, how nematode-associated microbiota (here including viruses) contribute to the assembly and functioning of a plant holobiont.

The objective of this study was to better understand the role of free-living nematodes in the stability of a plant holobiont. We hypothesized that (1) soil nematode communities and their vectored microbiota, specifically viruses, can increase the stability of a plant holobiont that is challenged by parasitic nematodes; (2) specific plant holobionts can condition the nematode community and their vectored microbiota, resulting in increased resistance of the plant holobiont that subsequently grow in the same soil. We use the resistance of the model plant holobiont tomato against invasion of the parasitic root-knot nematode *Meloidogyne hapla* as measure for the stability of the holobiont. The term resistance refers here to the ability of the plant holobiont to restrict infection of a pathogen or pest, so that the potential effects of disease are reduced. To test these hypotheses, we conducted a series of pot experiments. Each experiment comprised a phase for conditioning of the soil biota by different plant species, followed by the initiation and assembly of the tomato holobiont, and finally the testing of the response of this holobiont to the parasite as a function of the preceding conditioning of the soil biota (Fig. 1). In this holistic study, we did not analyze the taxonomic structure of the microbiome resulting from holobiont assembly. Our primary interest was in the holobiont’s resistance as conferred by functional networks, rather than the contribution of individual microbial species to the holobiont’s resistance.

**Fig. 1.**
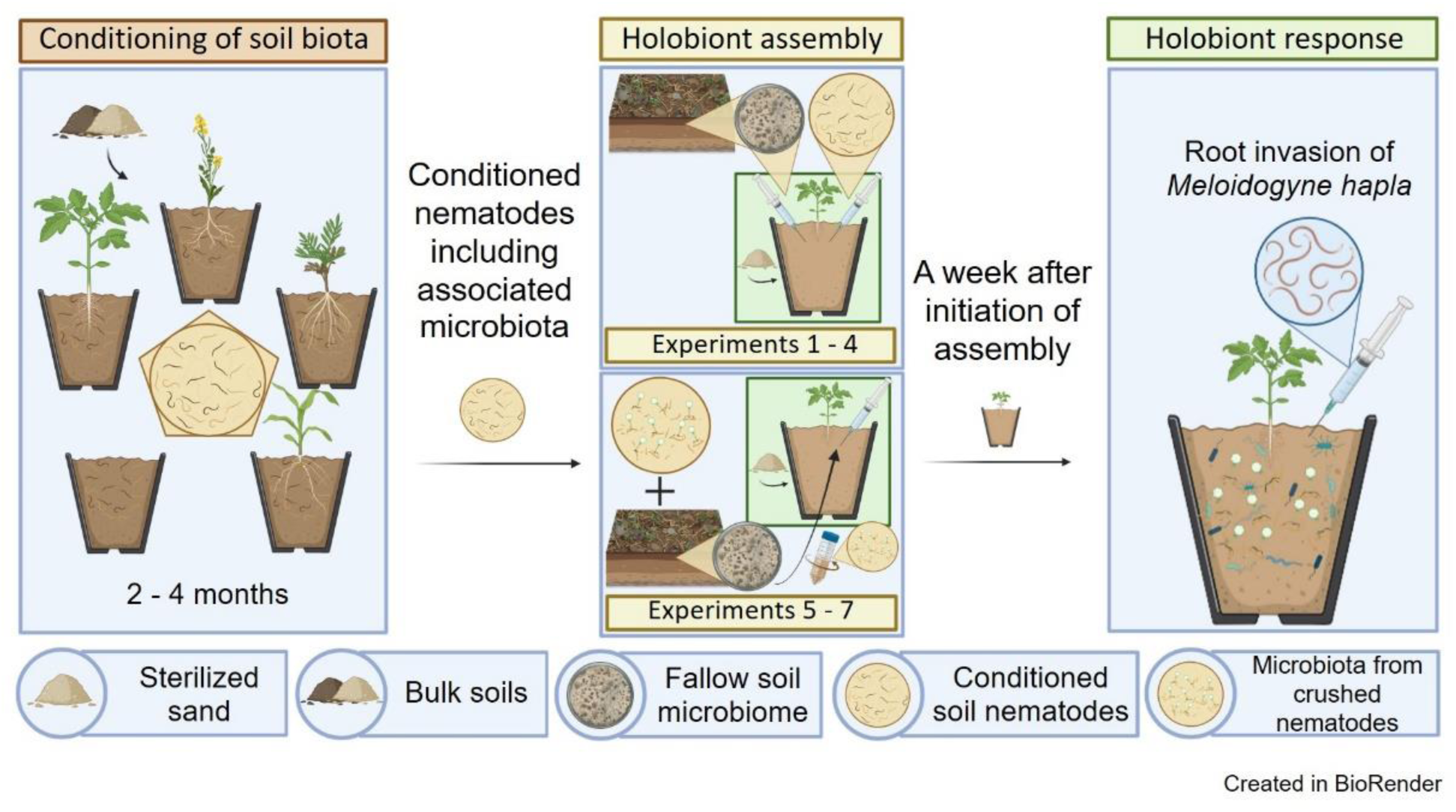
Conceptual figure of the study design. This study comprises a series of seven experiments, each of which is composed of three phases: 1) conditioning of the soil biota, including nematode communities, by plants; 2) initiation and assembly of the holobiont; and 3) response of the holobiont to a pathogen. During the conditioning phase, fallow arable soil with its natural biome was used to grow tomato (*Solanum lycopersicum* L.), oilseed rape (*Brassica napus* L.), marigold (*Tagetes patula*) or maize (*Zea mays* L.) for two to four months under controlled conditions, or the soil was left fallow. The conditioned nematode communities were extracted and combined with a nematode-free microbiome from fallow soil and a tomato seedling in order to initiate the assembly of the plant holobiont (Experiments 1–4). Alternatively, in Experiments 5-7, the conditioned nematodes were crushed to use only the nematode-associated microbiota for the initiation of the holobiont assembly. In the response phase, the assembled tomato holobionts were challenged with the root-invading nematode *Meloidogyne hapla*. The number of root-invaded nematodes was used as an indicator of holobiont resistance.

## MATERIALS AND METHODS

### Conditioning of soil nematode communities by plants

Soil samples harboring their native biota were collected from fallow plots of an arable field near Ahlum, Germany (52°10’00“N 10°35’00”E). One part of the field is managed conventionally while the other part is under certified organic farming since 2001 (according to EU Organic Guidelines, organic certification number DE-NI-039-00451-A), including the use of organic fertilizers, and pest control without pesticides. Depending on the crop, the organic field plot is tilled once a year in spring or autumn, and harrowed two to three times and/or hoed one to two times per year using a roller or tine hoe. The conventional field plot is subject to reduced tillage. Extended crop rotations of at least five years were similar for both parts of the field. The soil was sandy loam with 1% humus and a pH of 7.1. For each experiment, eight to ten fresh soil samples were collected from separate spots within a 20 m² area, corresponding to eight to ten replicates of the experiment. The soils were mixed with 50% sterile sand to prevent the pots from clogging. Seeds were planted in two-liter pots filled with this soil mixture. Plant varieties known to be robust against typical pathogens were used to keep the plants healthy during the conditioning phase. These varieties included tomato (*Solanum lycopersicum* L.) cultivar ’Bellandine F1’ (Kiepenkerl, Everswinkel, Germany), oilseed rape (*Brassica napus* L.) cultivar ’LG Adonis’ (Limagrain, Edemissen, Germany), marigold (*Tagetes patula*) cultivar ’Petite Orange’ (Sperli, Everswinkel, Germany), and maize (*Zea mays* L.) cultivar ‘SY Liberty’ (Syngenta Agro, Frankfurt am Main, Germany). The pots were arranged in a Randomized Complete Block Design. The plants were cultivated for two to four months, either outside or in the greenhouse, depending on the season. In some of the experiments, control pots were left fallow and treated like planted pots, but with less watering. Greenhouse conditions were 24 °C and a 16/8 h photoperiod, fertilization was not needed.

At the end of the conditioning phase, the nematodes from 200 ml of soil from each pot were extracted by centrifugal floatation using MgSO_4_ at 1.18 specific density (Hooper et al. 2005). The nematodes were collected from the float on a 20-μm sieve, washed with sterile water, and transferred with 50 ml sterile water to a glass beaker. These conditioned nematode communities were then used to either initiate plant holobiont assembly or investigate nematode-associated microbiota.

### Retrieving microbiomes from fallow soil to initiate holobiont assembly

At the end of the conditioning phase, fresh fallow soil was collected from the arable field near Ahlum. The soil was 2-mm sieved and suspended in three volumes of water in a large tray that was shaken horizontally overnight at 20 °C. The soil suspension was decanted into one-liter centrifuge tubes to pellet most of the soil particles at 500 *g* for five minutes. The supernatant was passed through 100-µm and 20-µm sieves to remove indigenous nematodes. This suspension of the fallow soil microbiome was used as a resource for holobiont assembly (see the respective paragraph below).

### Preparation of nematode-associated microbiota

To prepare a suspension of nematode-associated microbiota, each conditioned nematode community was pelleted by centrifugation at 4000 *g* for 10 min at 4 °C, and transferred with 500 µl phage-buffer (10 mM Tris-Cl, 10 mM MgSO_4_, pH 7.0) to a mortar. The nematodes were mechanically homogenized with a sterile pestle and then suspended in 5 ml phage-buffer. An aliquot of each suspension was mounted on a glass slide and examined on an SZX12 stereo microscope (Olympus, Tokyo, Japan) to confirm the absence of intact nematodes. To check for phages, a 25 µl aliquot of each suspension was pipetted onto parafilm. A 400-mesh copper grid was placed on the droplet for 5 min. The copper grids had been coated with Pioloform and stabilized with carbon before usage in virus detection. After washing with water to remove unbound material, the grid was stained with 1% uranyl acetate in Millipore water and examined at 80 kV using a transmission electron microscope (TEM). Images were taken at 42,000 x or 67,000 x magnification using a 2080 by 2080 pixel charge-coupled device sensor camera “Veleta” (Olympus), side mounted to the TEM column.

Each suspension of nematode-associated microbiota was divided into two tubes. One of them was heated to 70 °C in a water bath for 10 min to inactivate microbiota, which served as a control, while the other was left untreated (active microbiota).

### Treatment of soil microbiome by nematode-associated microbiota prior to holobiont assembly

Twenty ml of the fallow soil microbiome suspension were inoculated in 50-ml tubes with 2 ml of the nematode-associated microbiota, either active or inactivated. In the sixth experiment (Fig. 1), an additional control with non-crushed nematodes was included. The tubes were shaken horizontally at room temperature for 48 hours to allow the active phages from the nematode-associated microbiota to encounter and infect their bacterial hosts. This mimics vectoring by nematodes. At the 24-hour mark, each tube was briefly opened under a laminar flow hood for five minutes to allow air exchange in a sterile environment.

After treatment, serial dilutions of the soil microbiome suspensions were plated on R2A medium supplemented with cycloheximide (100 µg/ml) to determine total bacterial CFU, and on King’s B medium supplemented with cycloheximide (100 µg/ml), chloramphenicol (13 µg/ml), and ampicillin (100 µg/ml) to determine CFU of pseudomonads after four days at 28 °C.

### Assembly of the holobiont

The surface of tomato seeds cultivar “Moneymaker” (Kiepenkerl, Everswinkel, Germany) was disinfected by sequentially washing three times with autoclaved tap water, once with 70% ethanol, once with 1.5% sodium hypochlorite, and four times with autoclaved tap water. Seeds germinated in 50-ml pots filled with wet sterile quartz sand (0-1 mm, Bötel-Mascheroder GmbH, Braunschweig, Germany). In the first four experiments (Fig. 1), two-week-old seedlings were inoculated with the fallow soil microbiome by drenching 50 ml of the suspension close to the stems. In treatments without microbiome (Experiment 2 and 3), seedlings received only water. The following day, the various nematode fractions were added to the pots by inoculating four holes made with a dibber in the sand around the stem with the respective suspension (4 x 4 ml). In the fifth, sixth and seventh experiment (Fig. 1), which involved nematode-associated microbiota, the fallow soil microbiome was inoculated to the pots after microbiota treatment by drenching 25 ml of the suspension close to the stems. The plants were then incubated in the greenhouse for one week to allow the rhizobiome to establish, before the root invasion assay. The greenhouse conditions were 24 °C, a 16/8 h photoperiod and pots were watered daily as needed.

### Assay of root invasion by parasites as a measure of holobiont resistance

*M. hapla* was reproduced on tomato cv. Moneymaker in the greenhouse. Eggs were extracted from galls by blending the roots in 1.5% sodium hypochlorite for 30 s and collecting the eggs on a 20-μm sieve (Bezooijen 2006). The egg suspension was transferred on a modified Baermann tray (Hooper et al. 2005) and freshly hatched J2 were collected. Before use, J2 were purified by allowing them to move through paper tissue to the bottom of a modified Baermann funnel. Each root system was challenged with 500 second stage juveniles of *M. hapla*. After one week, roots were harvested. For microscopic counting, the root-knot nematodes in the roots were stained by acid fuchsin (Bybd et al. 1983). The roots of each plant were incubated in 1.5% sodium hypochlorite for 3 minutes for clearing, then rinsed and soaked in tap water for 15 minutes, and stained in 40 ml of 1% acid fuchsin solution. The beakers were heated in a microwave until near-boiling and then cooled. Roots were rinsed under running tap water, transferred to 20 ml of acidified glycerin, and stored at 4 °C prior to microscopic counting of root-knot nematodes using an Olympus SZX12 stereo microscope at 90 x magnification.

In the fifth experiment, we investigated the effect of the microbiome treatment on the defense response of the tomato plants to root-invading parasites. Leaf samples of each tomato plant were immediately frozen in liquid nitrogen seven days after inoculation of *M. hapla*. RNA was extracted using a RNeasy kit (Qiagen, Hilden, Germany) according to the manufacturer’s protocol. RNA concentration was checked using nanodrop spectrophotometer and quality checked on 1.3% agarose gel. Reverse transcription and real-time PCR were performed using a Luna^®^ Universal One-Step RT-qPCR Kit (New England Biolabs, Frankfurt am Main, Germany) in 20 µl reactions containing 3 µl of RNA, Luna Universal One-Step Reaction Mix, Luna WarmStart RT Enzyme Mix, 10 µM of either ubiquitin primers GTGTGGGCTCACCTACGTTT and ACAATCCCAAGGGTTGTCAC (Bhattarai et al. 2008), or PR1A1 primers CTGGTGCTGTGAAGATGTGG and TGACCCTAGCACAACCAAGA (Harel et al. 2014). Thermocycles in a CFX Connect Real Time Detection System (Bio-Rad, Feldkirchen, Germany) were as follows: reverse transcription at 55 °C for 10 min, initial denaturation at 95 °C for 1 min, 40 cycles of a denaturation step at 95 °C for 10 s, an extension step at 60 °C for 30 s, and a step to read SYBR fluorescence at 80 °C for 5 s. Melt curves were analyzed to make sure that only the intended gene fragments contributed to the fluorescence at 80 °C.

### Statistical analyses

All experiments were done in Randomized Complete Block Design. Nematode counts in roots were compared by fitting generalized linear mixed models using the GLIMMIX procedure of the software package SAS 9.4 (SAS Institute Inc., Cary, NC), with Poisson distribution and applying a log link function. In experiments 1-3 with 2 x 2 design, the LSMEANS statement was used to estimate the effect slices with respect to the factors Soil or Microbiome, respectively (LSMEANS Conditioning * Soil / diff slice = Soil). Normal distribution was assumed for C_q_ data from quantitative PCR and log-transformed bacterial densities (CFU/ml). The Kenward–Roger procedure was applied to estimate data-dependent degrees of freedom to account for small sample numbers, resulting in more conservative tests. In experiments 4 and 7 with multiple comparisons, *P* values were adjusted by Tukey’s method using a LSMEANS statement (LSMEANS Treatment / adjust=Tukey ADJDFE=ROW plots=meanplot cl). Residual plots were used to check for model fit (plots=residualpanel). All boxplots were produced with R (R Core Team 2025) / RStudio (version 2026.01.0, Posit Software, PBC) using the ggplot2 package version 4.0.1 (Wickham 2016).

## RESULTS

### First experiment: Nematodes from tomato rhizosphere promoted resistance of the tomato holobiont more than those from oilseed rape

In a pot experiment in the greenhouse, nematode communities in arable soil with different management legacies were conditioned either by oilseed rape or tomato for four months. The soil with indigenous soil biota originated from either the conventionally or the organically managed part of an arable field. The conditioned nematode communities were inoculated to the rhizosphere of tomato seedlings that had received a 5 µm-sieved (nematode-free) microbiome from the respective fallow soil. After one week of rhizobiome assembly, the ability of the tomato holobionts to withstand invasion by root-knot nematodes was tested. For both soils, the conditioning of the nematode community by either oilseed rape or tomato significantly affected the number of nematodes invading tomato roots (Fig. 2). Nematode communities from tomato rhizosphere promoted the assembly of a more resistant tomato holobiont than those from oilseed rape (GLMM, factor Conditioning: *P* = 0.0001). The effect was more pronounced for the soil biota from the organically managed part of the field (GLMM, Conditioning*Soil: *P* = 0.0446, contrasts were *P* = 0.0109 for the conditioning effect in conventional soil and *P* = 0.0001 for organic soil). Management legacy of the arable soil had a significant effect, with biota from the conventionally managed area of the field conferring tomato a higher resistance to root-knot nematodes than those from the organically managed area (GLMM, factor Soil: *P* = 0.0001). In conclusion, the pre-crop had an effect on the nematodes and/or their associated microbiota, which affected the resistance level of tomato against parasitic nematodes in the presence of fallow soil microbiota. For the further experiments, we used the organically managed soil harboring the more responsive microbiota.

**Fig. 2.**
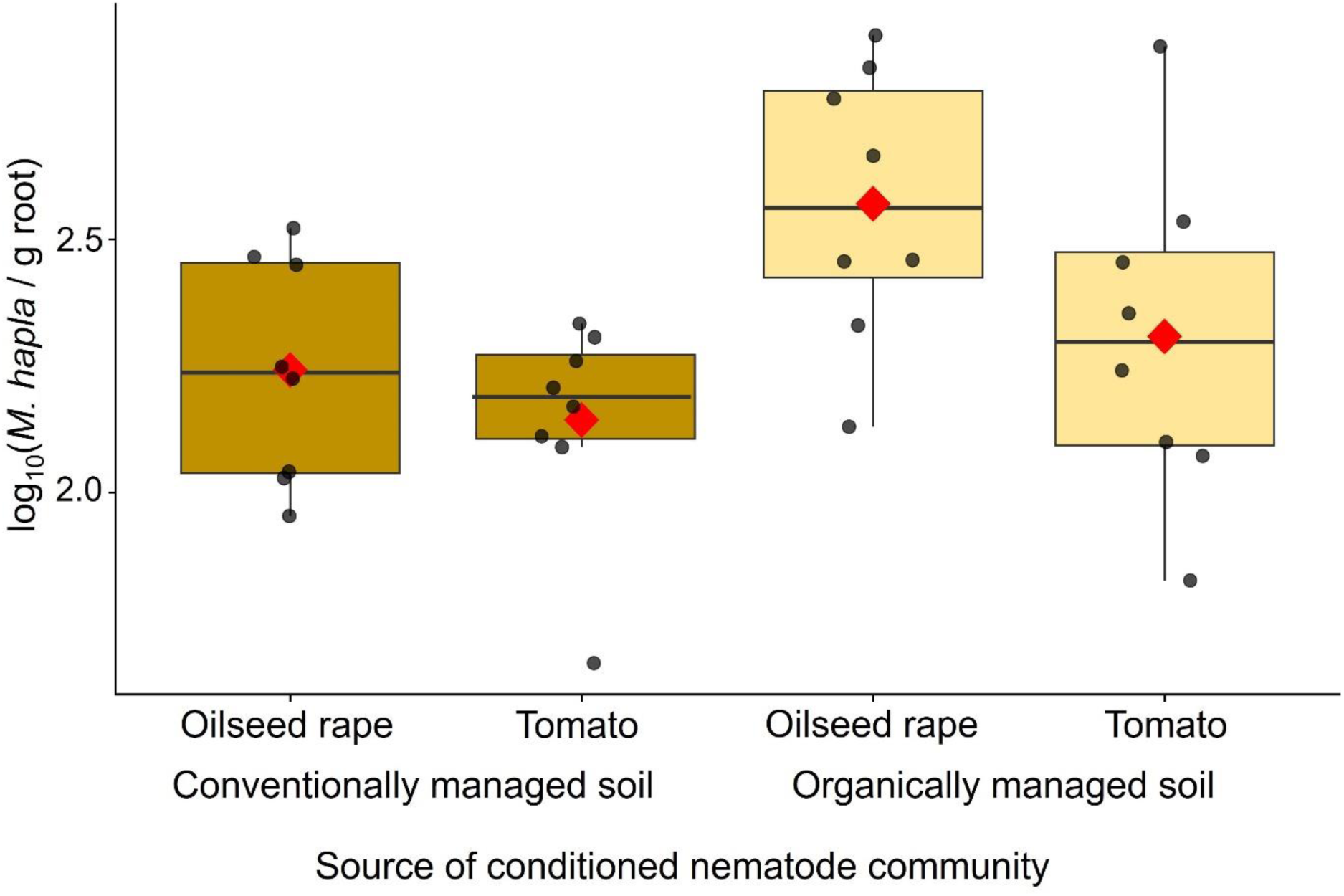
Effect of nematode communities conditioned by either oilseed rape or tomato on the tomato rhizobiome in terms of its suppression of root invasion by the parasite *Meloidogyne hapla* (Experiment 1). Soil from either the conventionally or the organically managed part of an arable field was used to condition the nematode communities in the rhizosphere of either oilseed rape or tomato. Assembly of the holobionts were initiated by inoculation of nematode-free fallow soil microbiome to pots with tomato seedlings growing in sterile sand, and inoculation of the conditioned nematode communities (including associated microbiota). A week later, the seedlings were challenged with *M. hapla*. Boxplots indicate the number of *M. hapla* that invaded the roots as a measure of the holobiont’s resistance. Each box spans the interquartile range of log_10_-transformed counts per g root; the horizontal line inside each box marks the median; the solid red diamond indicates the mean (N = 8).

### Second experiment: Nematodes from tomato rhizosphere promoted resistance of the tomato holobiont more than nematodes from fallow soil

Nematode communities were conditioned by growing tomato for three months in pots with an arable soil harboring indigenous nematodes. Pots without plants were treated in the same way to generate unconditioned nematode communities. The nematode communities were inoculated to pots with tomato seedlings that either received the nematode-free microbiome from the fallow soil or not. After one week, the resistance of the assembled holobionts to invasion by root-knot nematodes was tested. Tomato plants with inoculated microbiome were more resistant than plants that only received the nematode communities (Fig. 3; GLMM, factor Microbiome: *P* = 0.0001). Nematodes from the tomato rhizosphere promoted the formation of a more resistant tomato holobiont than those from the fallow soil in the presence of the nematode-free fallow soil microbiome (GLMM, factor Conditioning*Microbiome, effect slice “No Microbiome”: *P* = 0.0152), while this effect was not apparent without inoculated fallow soil microbiome (effect slice “With Microbiome”: *P* = 0.2489). In conclusion, the effect of conditioning the nematodes and/or their associated microbiota on the resistance level of tomato against parasitic nematodes depended on the presence of the soil microbiome in the assembly phase of the holobiont.

**Fig. 3.**
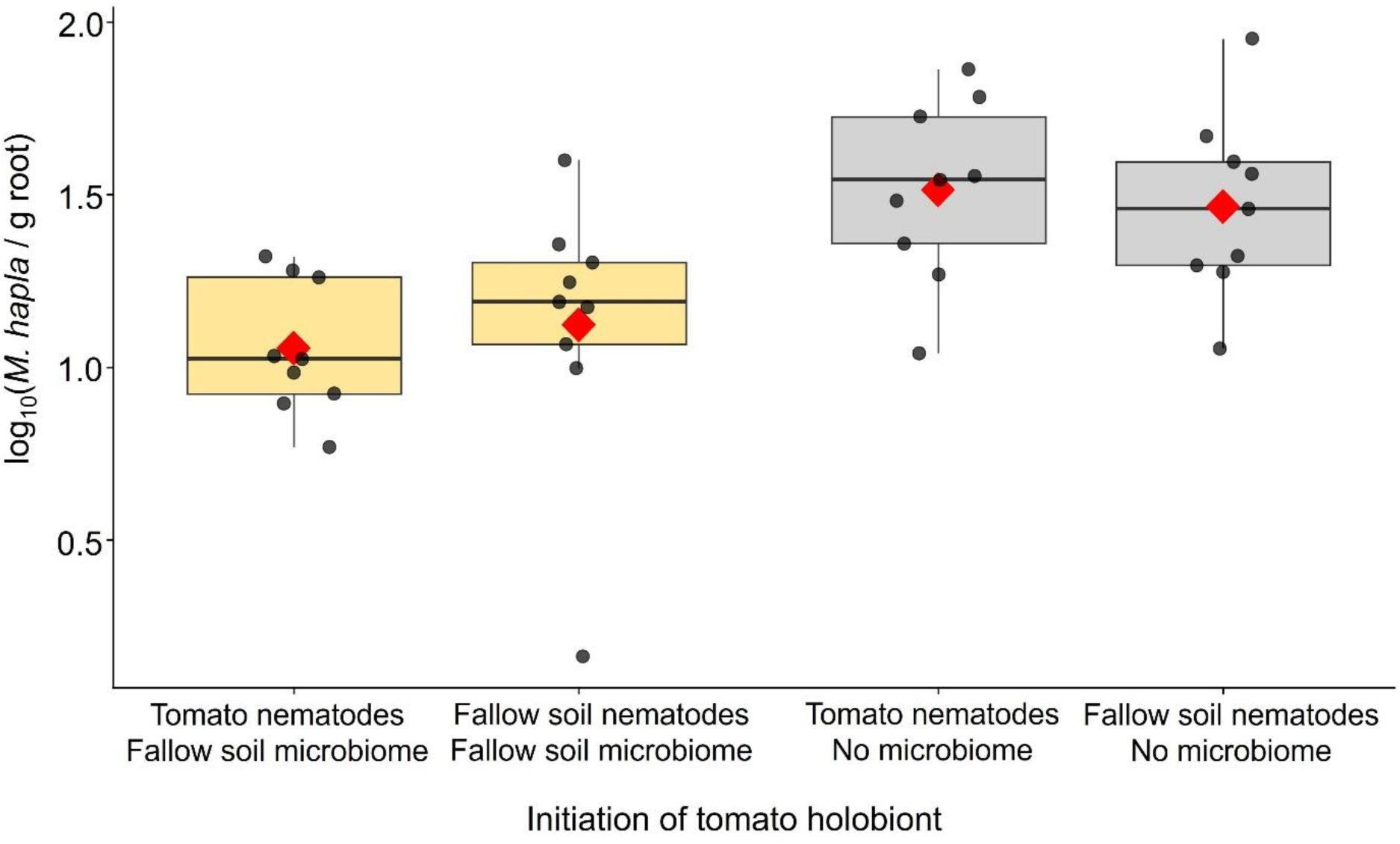
Effect of nematodes communities conditioned by tomato compared to unconditioned soil nematodes on the assembly of the tomato rhizosphere microbiome in terms of its suppression of root invasion by the parasite *Meloidogyne hapla* (Experiment 2). Fallow soil from an arable field was used to condition the indigenous nematode communities with or without tomato. The fallow soil microbiome and nematode communities were inoculated to roots of tomato seedlings for holobiont assembly. As controls for impaired holobiont assembly, nematodes that were either conditioned or not conditioned were added to tomato seedlings that had not been inoculated with a microbiome. A week later, the seedlings were challenged with *M. hapla*. Boxplots indicate the number of *M. hapla* that invaded the roots as a measure of the holobiont’s resistance. Each box spans the interquartile range of log_10_-transformed counts per g root; the horizontal line inside each box marks the median; the solid red diamond indicates the mean (N = 9).

### Third experiment: Effect of nematode communities conditioned by either tomato or tagetes on resistance of the tomato holobiont

Tagetes roots emit bioactive compounds that differentially antagonize nematodes, depending on exposure and susceptibility of the species. Therefore, we assumed a strong effect on nematode community structure in the rhizosphere during the conditioning phase. We compared the effect of soil nematode communities conditioned by either tomato or tagetes on resistance of the tomato holobiont to root invasion by the parasite *M. hapla*. Tomato holobionts that were assembled under the influence of tagetes-conditioned nematode communities showed significantly less root invasion by *M. hapla* than those modulated by tomato-conditioned nematodes (Fig. 4; GLMM, factor Conditioning: *P* = 0.0001). Presence of the fallow soil microbiome at the initiation of the tomato holobiont had a significant effect on later root invasion by *M. hapla* (GLMM, factor Microbiome: *P* = 0.0438). Interestingly, tomato inoculation with the tagetes-conditioned nematodes resulted in a lower root invasion than tomato-conditioned nematodes, with or without added fallow soil microbiome (GLMM, factor Conditioning*Microbiome, effect slice “With Microbiome”: *P* = 0.0074, effect slice “No Microbiome”: *P* = 0.0001). This suggests that the tagetes-conditioned nematodes or their associated microbiota antagonized root invasion by *M. hapla* directly, or via a plant-mediated mechanism.

**Fig. 4.**
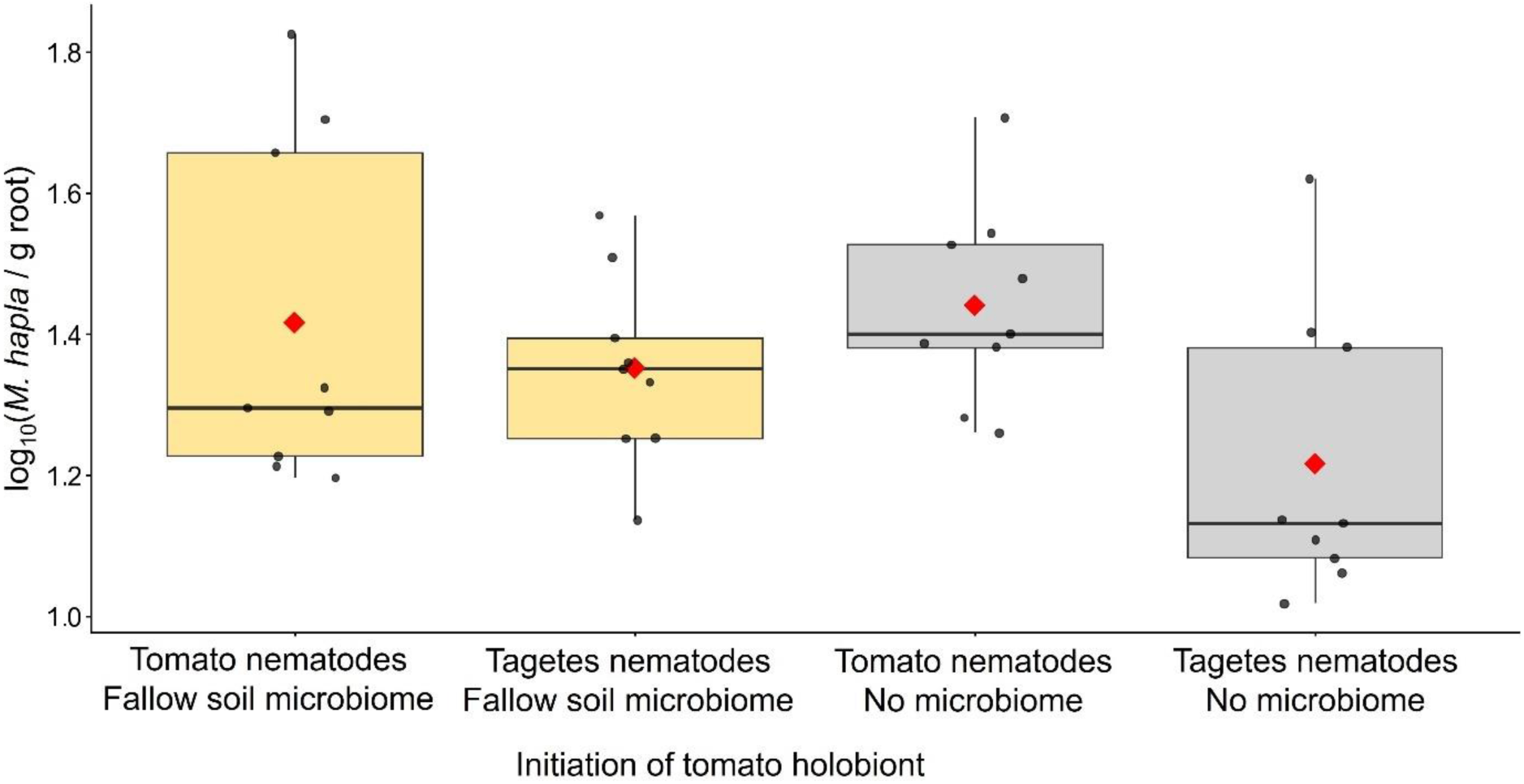
Effect of nematode communities conditioned by either tomato or tagetes on the assembly of the tomato rhizobiome in terms of its suppression of root invasion by the parasite *Meloidogyne hapla* (Experiment 3). Soil from the organically managed part of an arable field was used to condition the nematode communities either by tomato or tagetes. Fallow soil microbiome and nematode communities were inoculated to roots of tomato seedlings for holobiont assembly. As controls for impaired holobiont assembly, the conditioned nematode communities were added to tomato seedlings that had not been inoculated with a microbiome. A week later, the seedlings were challenged with *M. hapla*. Boxplots indicate the number of *M. hapla* that invaded the roots as a measure of the holobiont’s resistance. Each box spans the interquartile range of log_10_-transformed counts per g root; the horizontal line inside each box marks the median; the solid red diamond indicates the mean (N = 9).

### Fourth experiment: Effect of nematode communities conditioned by tomato, tagetes, or unconditioned on resistance of the tomato holobiont

We compared the effect of unconditioned fallow soil nematode communities to those conditioned by either tomato or tagetes on resistance of the resulting tomato holobiont to root invasion by the parasite *M. hapla*. The counts of parasitic nematodes in the root were significantly lower for tomato holobionts influenced by the plant-conditioned nematode communities in comparison to those influenced by unconditioned nematode communities (Fig. 5). Like in the third experiment, there was a trend of slightly lower parasite counts for the tagetes treatment compared to the tomato treatment, which was statistically not significant in this experiment.

**Fig. 5.**
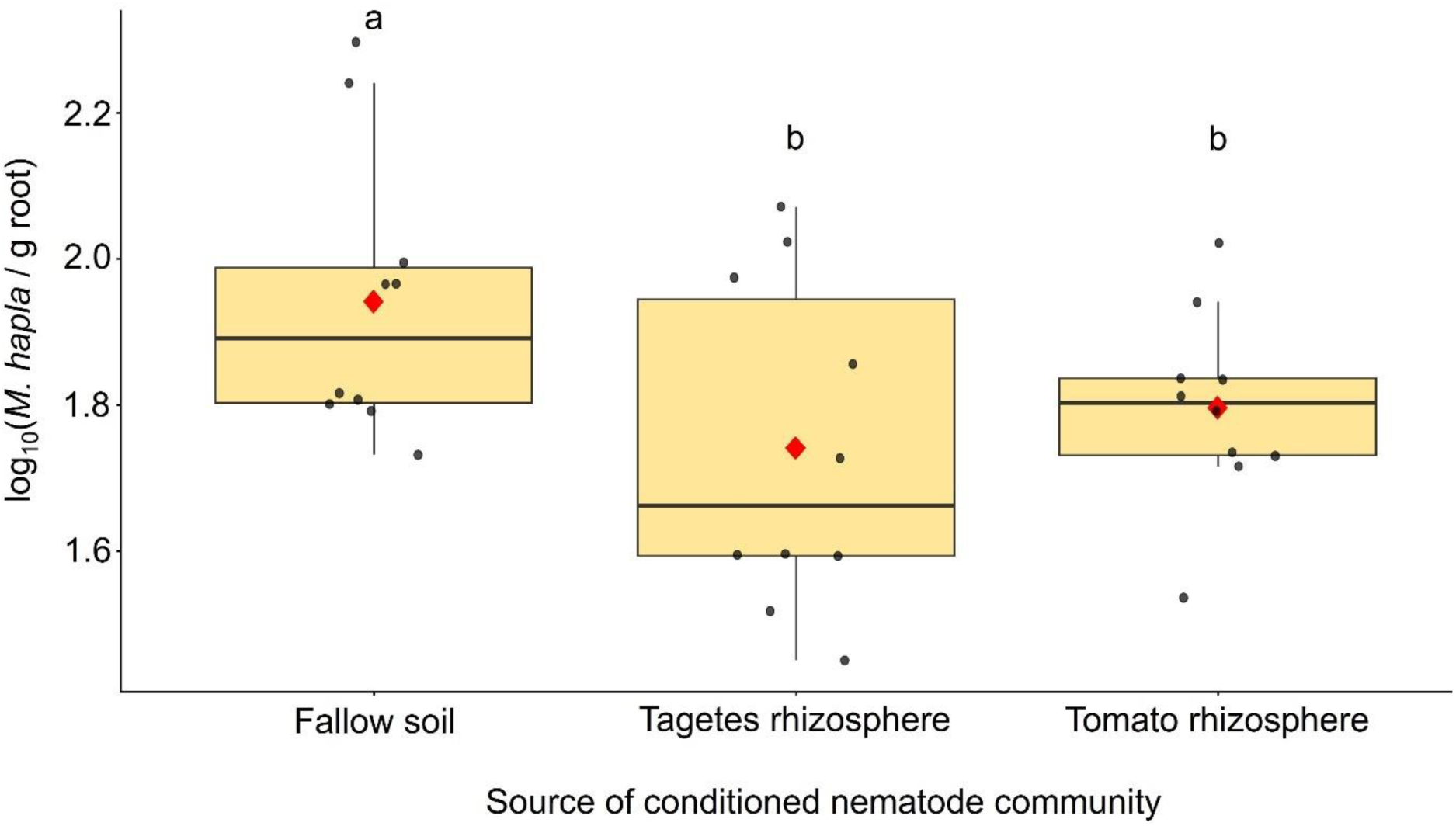
Effect of nematode communities conditioned by tomato or tagetes compared to unconditioned soil on the assembly of the tomato rhizobiome in terms of its suppression of root invasion by the parasite *Meloidogyne hapla* (Experiment 4). Soil from the organically managed part of an arable field was used either to condition the nematode communities by tomato or tagetes, or to retrieve an unconditioned nematode community. Fallow soil microbiomes and nematode communities were inoculated to roots of tomato seedlings for holobiont assembly. Boxplots show the number of subsequently inoculated *M. hapla* juveniles (J2) per root weight that invaded the roots as a measure of the holobiont’s resistance to the parasite. The box spans the interquartile range of log_10_-transformed counts; the horizontal line inside each box marks the median; and the solid red diamond indicates the mean. Statistical significance is indicated above the boxes (Tukey-adjusted *P* < 0.05, GLM with Poisson distribution, N = 10).

### Fifth experiment: Inclusion of microbiota from crushed nematodes in holobiont assembly increased resistance of tomato

We investigated whether the effects on holobiont resistance that were observed in the previous experiments, were caused by nematodes directly, or whether nematode-associated organisms like phages are contributing to the holobiont assembly. Tomato-conditioned nematodes were inactivated by crushing to obtain a suspension of microbiota that were associated with the nematodes. A microbiome suspension from the respective unplanted soil was treated with these microbiota, or a heat-inactivated control, and inoculated to the roots of tomato seedlings to initiate holobiont assembly. The parasite *M. hapla* was found to be less successful in invading the roots when the microbiota from nematodes contributed to the assembly of the holobiont, in comparison to the control group with inactivated microbiota (Fig. 6). Bacterial densities in the soil microbiome suspensions treated with active or inactivated nematode-associated microbiota before inoculation to tomato roots did not significantly differ, as determined by plating on King’s B media for pseudomonads (Fig. 7A), and R2A for total bacterial colony forming units (Fig. 7B). In line with the enhanced resistance of tomato holobionts influenced by nematode-associated microbiota, the plants showed a higher expression (i.e. lower Cq) of the defense gene PR1A1 in response to *M. hapla* compared to the control group (Fig. 7C, GLM, *P* = 0.0001). In the same RNA-extracts, transcript levels of the housekeeping gene ubiquitin did not significantly differ between both treatments. Using transmission electron microscopy, intact phages were detected in the suspension of the crushed nematodes, but not after heating to 70 °C (Fig. 7D).

**Fig. 6.**
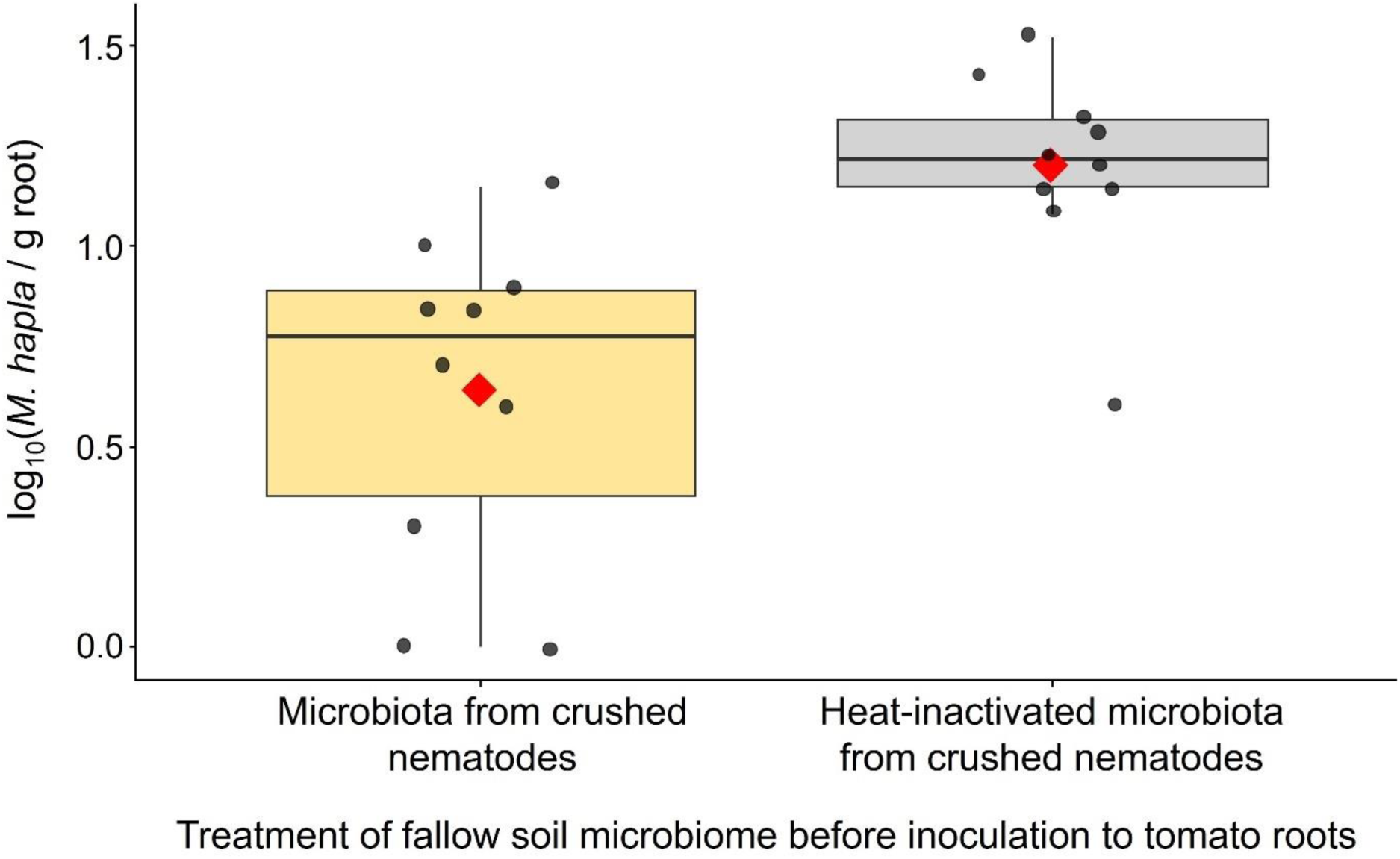
Effect of the inclusion of a microbiota from tomato-conditioned crushed nematodes in the assembly of the tomato holobiont on its resistance (Experiment 5). A fallow soil microbiome suspension was inoculated with microbiota from crushed nematodes that originated from the rhizosphere of tomatoes. In the control, the microbiota were heat-inactivated (70 °C). After two days, the microbial suspensions were inoculated to the roots of tomato seedlings to initiate holobiont assembly. Subsequently, *M. hapla* juveniles were inoculated to the roots. Boxplots show the number of *M. hapla* in the roots. The box spans the interquartile range of log_10_-transformed counts; the horizontal line inside each box marks the median; the solid red diamond indicates the mean (N = 10).

**Fig. 7.**
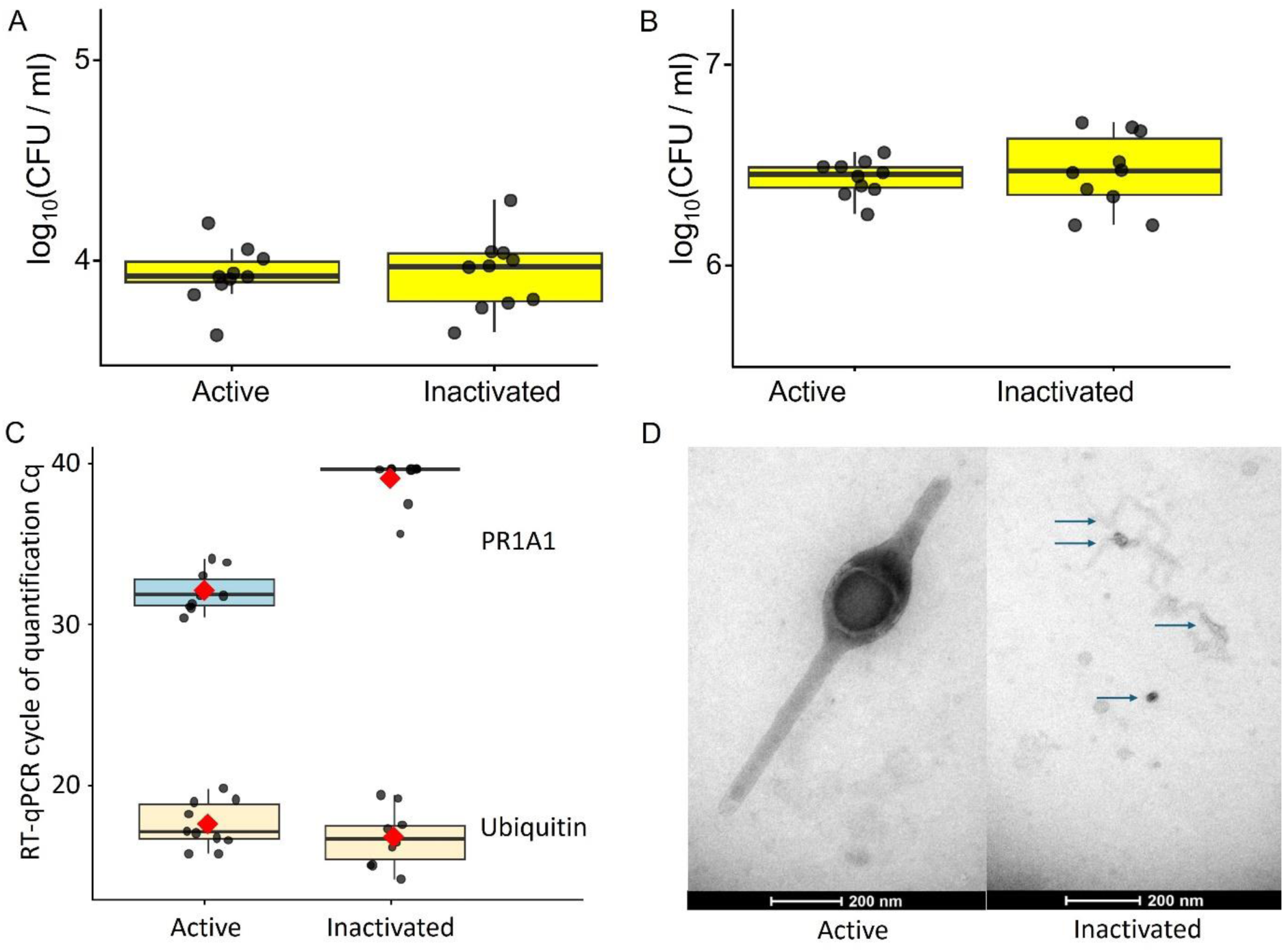
Microbiome, plant defense, and phages (Experiment 5). Bacterial densities in the soil microbiome suspensions treated with active or inactivated nematode-associated microbiota. Colony forming units (CFU) were counted on (A) R2A and (B) King’s B agar plates. (C) Reverse transcription quantitative real-time PCR of the tomato defense gene PR1A1 and the reference housekeeping gene ubiquitin in leaves 7 days after inoculation of *Meloidogyne hapla*. The box spans the interquartile range; the horizontal line inside each box marks the median; the solid red diamond indicates the mean (N = 10). No signal was treated as Cq = 40. (D) In the suspension of crushed nematodes, phage particles were detected by transmission electron microscopy. The left panel shows a bicaudavirid-like phage comprising head with two tails. The blue arrows in the lower right panel indicate phage fragments released from the capsid in the corresponding heat-inactivated suspension.

### Sixth experiment: Inclusion of 0.2 µm filtrates from crushed nematodes in holobiont assembly increased resistance of tomato

Suspensions of crushed tomato-conditioned nematodes were 0.2 µm-filtered to obtain a filtrate of nematode-associated phages. Microbiome suspensions from fallow soil were treated either with these phages, or the phage-free buffer. Microbiome suspensions were inoculated to the roots of tomato seedlings to initiate holobiont assembly. The parasite *M. hapla* was found to be less successful in invading the roots when the nematode-associated phages contributed to the assembly of the holobiont, in comparison to the control without nematode-associated phages (Fig. 8).

**Fig. 8.**
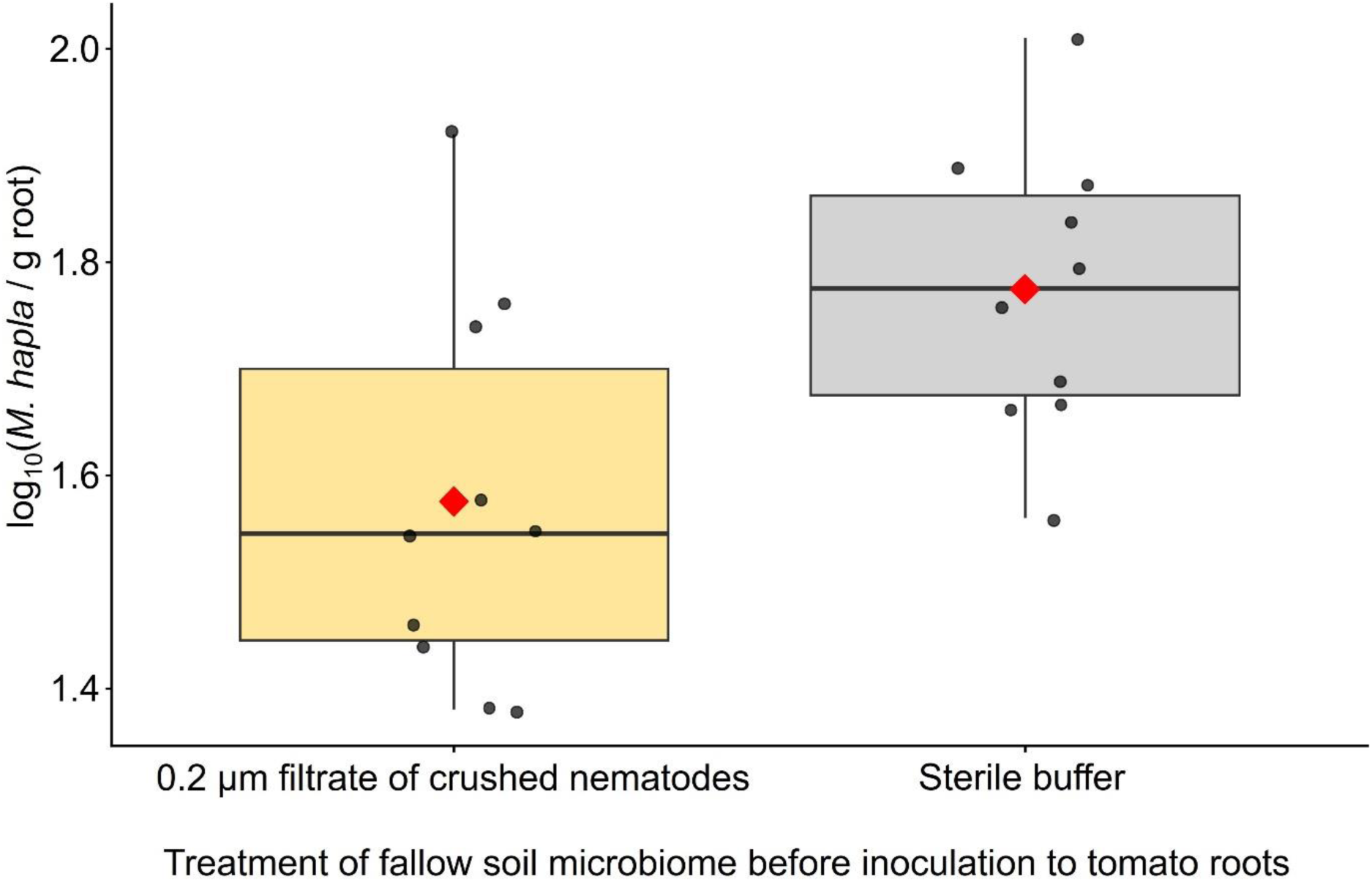
Effect of the inclusion of a sterile filtrate from tomato-conditioned crushed nematodes in the assembly of the tomato holobiont on its resistance (Experiment 6). A fallow soil microbiome suspension was shaken for two days either inoculated with a 0.2 µm filtrate from tomato-conditioned crushed nematodes, or with sterile buffer. The suspensions were drenched to the roots of tomato seedlings and incubated for one week to allow the rhizobiome to establish before the root invasion assay. After a week, *M. hapla* juveniles were inoculated to the roots. Boxplots show the number of *M. hapla* in the roots. The box spans the interquartile range of log_10_-transformed counts; the horizontal line inside each box marks the median; and the solid red diamond indicates the mean (N = 10).

### Seventh experiment: Phages associated with nematodes from tomato but not maize rhizosphere increased microbiome-mediated resistance of tomato

As already observed in experiment 5, *M. hapla* was less successful when the tomato holobiont was assembled starting from fallow soil microbiome and phages from crushed tomato-conditioned nematodes, compared to the control with heat inactivated phages (Fig. 9). However, when nematode communities were conditioned in maize rhizosphere in the same soil, then the nematode-associated phages did not modulate the soil microbiome in terms of reducing *M. hapla* counts in tomato roots. This suggests that the biota stabilizing the holobiont were plant-specific. Interestingly, intact tomato-conditioned nematodes harboring their associated microbiota (including phages) did not have an effect on *M. hapla* counts in tomato roots, suggesting that the microbiota attached to the nematodes were not mobile enough to interact with microbes in the fallow soil suspension.

**Fig. 9.**
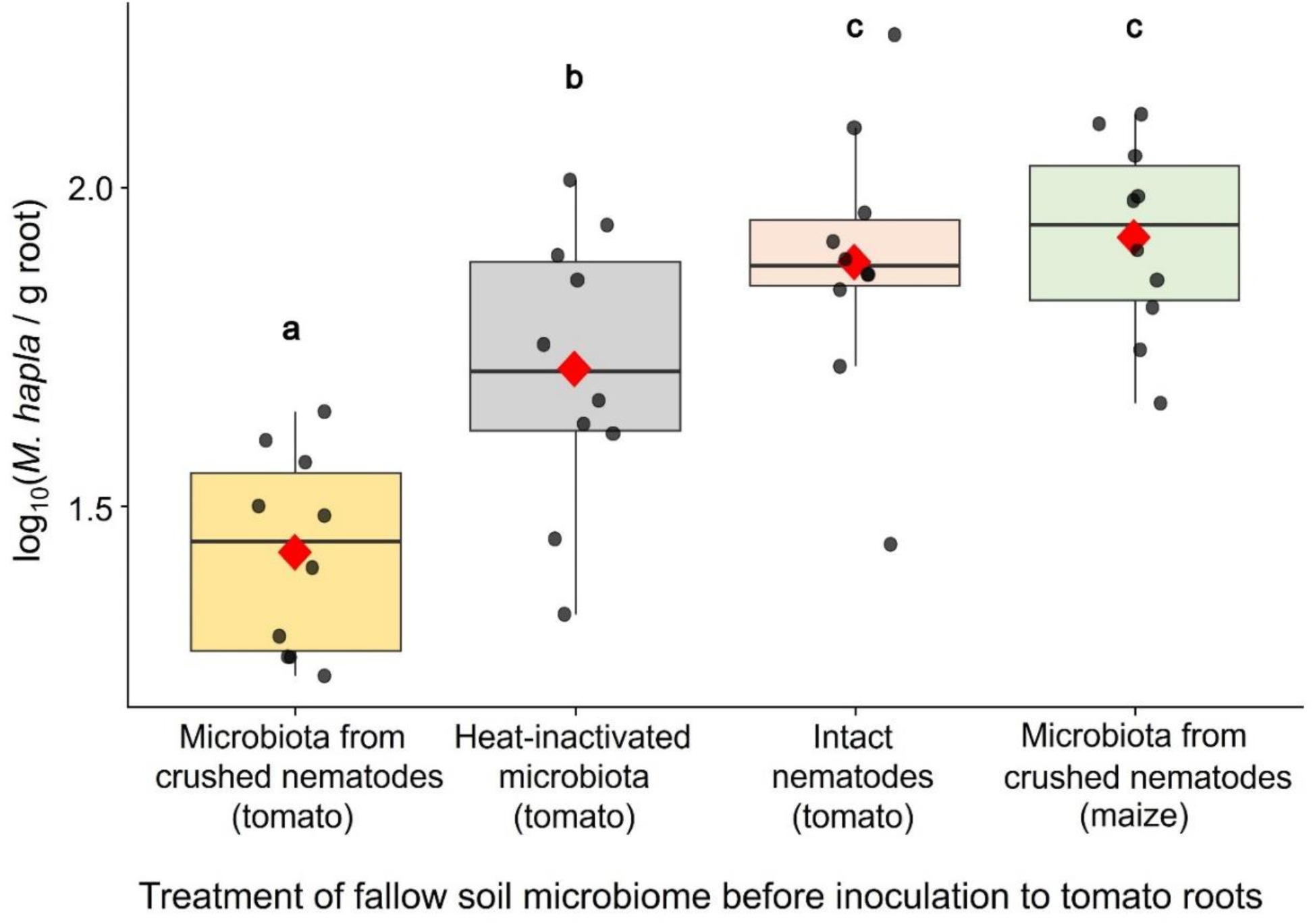
Effect of microbiota associated with nematodes from tomato or maize on assembly of the tomato holobiont in terms of its suppression of root invasion by the parasite *Meloidogyne hapla* (Experiment 7). A suspension of fallow soil microbiome was treated with crushed nematodes from the rhizosphere of either tomato or maize to allow nematode-associated phages to infect their bacterial hosts. In the controls, inactivated nematode-associated microbiota (70 °C) or intact nematodes (with associated active microbiota) were incubated with the microbiome suspension. The treated microbial suspensions were then inoculated to the roots of tomato seedlings for a week, before challenging the holobiont by inoculation of *M. hapla*. Boxplots show the log_10_-transformed number of J2 per gram root. Each box spans the interquartile range; the horizontal line inside each box marks the median; the solid red diamond indicates the mean. Statistically significant differences are indicated by different letters above the boxes (Tukey-adjusted *P* < 0.05, GLM with Poisson distribution, N = 10).

## DISCUSSION

This study indicates that nematode communities and their associated microbiota contributed to the self-organization of the tomato holobiont, as evidenced by increased resistance to root-knot nematodes. Preceding plant holobionts conditioned these biota, leading to a biotic legacy in soil. Our experiments comprised three phases. First, the nematode community and their associated microbiota were conditioned by a preceding plant in order to mimic biotic soil legacy. Second, holobiont assembly in the root zone of the target plant (tomato) was initiated in a sterilized pot system by the interaction between the conditioned nematodes and/or their associated microbiota and a fallow soil microbiome, under the influence of the root. Third, the assembled holobiont responded to invading parasites (*M. hapla*) through the concerted action of plant- and microbe-mediated defenses. These three phases are discussed in the following.

### Biotic legacy of preceding plants

We found that tomato and tagetes conditioned the fallow soil nematode community in a manner that this subsequently contributed to a more resistant tomato holobiont. Changes of the nematode community were plant-specific, persisted across plant generations and thereby structured the functional network of subsequent holobionts. Changes in soil nematode communities specific to a given plant have been well documented for root-feeding nematodes, but are less well studied for microbe-feeding nematodes (Yeates 1999). Plants with their specific root exudates and microbiomes provide the resources for microbe-feeding nematodes, leading to much larger populations in the rhizosphere than in fallow soil. Several studies have demonstrated that plant species identity has a strong effect on nematode community composition (Ikoyi et al. 2023; Deyn et al. 2004; Viketoft et al. 2009). The quality of plant resources, including specific root exudates, root architecture and plant-specific microbiomes, has been reported to be more important than the resource quantity for structuring nematode communities, but the underlying mechanisms are not well understood (Cortois et al. 2017).

In our study, tomato could better modulate the soil nematode community than oilseed rape in terms of the subsequent assembly of a resistant tomato holobiont. This implied that biota from foreign plants tend to be less beneficial (Elhady et al. 2018). Surprisingly, tagetes as preceding plant had an equal or even better effect on the beneficial potential of nematode communities to build a resistant tomato holobiont. Tagetes releases thiophene compounds such as α-terthienyl from the roots that antagonize diverse root-feeding and microbe-feeding nematodes, thereby strongly influencing nematode community structure in its rhizosphere (Kanfra et al. 2021; Hamaguchi et al. 2019). These bioactive compounds of tagetes were removed by washing the nematode community on a sieve, so they should not directly affect the subsequent tomato holobiont assembly. Indirectly, the structure of the nematode community and thereby the phages and microbes associated with these nematodes could have been influenced by bioactive compounds released from tagetes roots. In the third experiment, this nematode fraction alone induced a more resistant tomato holobiont than the nematode-mediated modulation of the soil microbiome. This was in contrast to the nematode fraction conditioned by tomato, that did not support the assembly of a resistant tomato holobiont without addition of a soil microbiome (first experiment). The nematode fraction includes specific nematode-associated microbiota (Elhady et al. 2017), which might be responsible for the differential effect of the nematode fractions from the tomato and tagetes rhizospheres. In this study, nematode-associated microbiota (including phages) were identified as an important legacy of preceding plant holobionts that persist across plant generations and thereby structure the functional network of subsequent holobionts.

Agronomically, effects of preceding crops on the yield of a following crop were previously shown (Jacobs et al. 2018). Hu et al. (2025) investigated the influence of different crop rotation schemes in a long-term field experiment on the maize rhizobiome and its effect on maize performance. Transplanting of the rhizobiome from a wheat-soybean-maize rotation scheme to maize seedlings resulted in enhanced plant performance, compared to rhizobiomes from less diverse wheat-maize rotation schemes. Ecologically, our results imply that soil nematodes and their associated microbiota (incl. phages) contribute to the temporal memory of ecosystems by encoding plant legacies. Such biotic memory can affect colonization of plants subsequently growing in the same soil by ecological filtering. From an applied perspective, crop rotation schemes and soil management practices that alter nematode–microbiota networks could be leveraged to steer plant holobiont assembly towards beneficial states. It needs more studies to understand the effects of organic farming on the responsiveness of crop holobionts to preceding plants, as indicated in the second experiment.

### Holobiont assembly

In contrast to the plant-centric view of a plant-governed assembly of plant-associated microbiota, the holobiont concept emphasizes the self-organization of the local microbiota through interactions among each other and with the plant. Our results extend current concepts of plant holobionts by identifying nematode communities and their associated microbiota as vital biotic components that act as ecological filters in the assembly of the plant holobiont. This ecological filtering involves selective feeding by microbivorous nematodes. It has been shown that bacteria-feeding nematodes affect bacterial community composition (Griffiths et al. 1999; Blanc et al. 2006; Postma-Blaauw et al. 2005). Similarly, fungi-feeding nematodes have pronounced prey-selecting preferences and will consequently alter the structure of fungal communities in the rhizosphere (Ruess et al. 2000). Modulation of the microbiome structure may therefore depend on the species composition of the conditioned nematode community. Moreover, bacteria-feeding nematodes were shown to adapt their feeding preferences based on epigenetically inherited learned avoidance of nematode pathogens (Kaletsky et al. 2025; Sengupta et al. 2024). Given our study’s holistic approach, it remains hypothetical whether conditioning nematodes and their associated microbiota led to changes in the taxonomic structure of holobiont assemblies. The functional network of species that constitutes the plant holobiont is defined by the interactions between its members. These interactions have recently been shown to influence the network members’ proteomes more than abiotic conditions (Moraïs et al., 2026). Microbial interactions often depend on direct contact or close proximity, and local niches (Jeckel et al. 2023; Bourceau et al. 2023). This is particularly true for cooperation, competition, metabolic coupling, and signal transduction, as these processes decline sharply over greater distances. Therefore, spatial structure may play a more significant role for functional networks than taxonomic microbiome structure. At a given point in time, biotic interactions may only be realized by some of the species present and in close proximity to their interaction partners. Moreover, the relative abundance of many species may just react to a changed holobiont without contributing to its functional network (Li et al. 2023). Consequently, the taxonomic structure of the total microbial community may not well reflect the structure of the functional network. Functional networks may be rather small. La et al. (2024) demonstrated that root-knot nematode invasion was lowest when communities comprising six or more native bacterial strains were present in the rhizosphere; single antagonists were unable to provide this degree of protection.

Soil nematodes carry specific microbiota, depending on the nematode species and biotic environment (Elhady et al. 2017; Topalović et al. 2022; Topalović et al. 2019). Recently, van Sluijs et al. (2025) demonstrated that bacteria-feeding nematodes (*Caenorhabditis* spp.) vector phages on their cuticle or after ingestion. Thereby, the investigated phage Φ Ppu-W11 was transported through soil to infect its host, *Pseudomonas putida*. Phages are shaping microbiomes in soil and other habitats by specifically lysing their hosts and through other mechanisms (Liang et al. 2025; Batinovic et al. 2019). In our experiments, the total phage community, together with other microbes, was released from the nematode fraction by crushing the nematodes. A fallow soil microbiome was exposed to the released microbiota under constant shaking, mimicking the vectoring of phages by nematodes. Initiation of the holobiont assembly by inoculation of the treated microbiome to tomato roots affected the resistance of the holobiont, depending on the pre-plant that conditioned the nematode-associated microbiota. We successfully applied TEM to qualitatively detect phages released from nematodes by crushing. The effective microbiota were most likely phages, because they multiply fast in the presence of their host and have the potential to change the bacterial community in the fallow soil microbiome suspension. In contrast, bacteria attach only in low numbers to nematodes (Elhady et al. 2021b), relative to their number in the soil suspension (Fig. 7). Bacteria and fungi have probably not the potential to sufficiently propagate and affect the fallow soil microbiome suspension within the two day incubation period. Intact nematodes including their associated microbiota were included in the seventh experiment as a control treatment. Their attached microbiota had no effect on the ability of the fallow soil microbiome to suppress *M. hapla*. This may be explained by the lack of mobility of soil nematodes in suspension. In soil, they move within pores with a diameter of 25–100 μm, but are immobile when such surface structures are lacking (Neher et al. 2019). It appears that the microbiota were not released from the intact nematodes, but were instead tightly attached to them. The mode of attachment of microbes or phages to nematode surfaces is not completely understood (Davies et al. 2023).

In accordance with our study on nematodes and their associated microbiota, a top-down control of plant microbiota for enhanced performance was also observed for predatory protists (Guo et al. 2024a; Guo et al. 2024b). This emphasizes the importance of self-organization of functional networks for the assembly of the plant holobiont, to which the bottom-up control by the plant contributes. Similar to our study, Kato et al. (2026) showed in a pot experiment a contribution of microbe-feeding nematodes to suppression of the root-knot nematode *Meloidogyne incognita* on green pepper. Densities of the parasite were significantly lower in pots with microbe-feeding nematodes than in those with microbes only, and the bacterial community structure differed.

### Response of the plant holobiont to invading parasites

We used the term resistance here as the ability of the plant holobiont to restrict invasion of a pathogen. In our study, resistance to invasion by the endoparasitic root-knot nematode *M. hapla* depended on the biotic resources for initiation of the tomato holobiont. This is consistent with previous research indicating that the rhizobiome provides basic protection against plant-parasitic nematodes, depending on the conditioning of soil microbiota by pre-crops (Topalović et al. 2020a; Elhady et al. 2021b; Elhady et al. 2025a; Elhady et al. 2025b). Various mechanisms are involved in this defense, some of which are based on the attachment of specific microbes to the nematode cuticle (Topalović et al. 2020b). La et al. (2024) demonstrated that simplified synthetic communities of root-associated bacteria could protect cucumbers from root-knot nematodes. The antagonistic mechanisms of the bacteria included hindrance of chemotaxis, suppression of migratory activities, secretion of anti-nematode substances, and modulation of plant defense responses. In our study, evidence for the latter mechanism was shown by upregulation of the defense gene PR1A1 in the microbiota-modulated tomato holobiont in the fifth experiment. La et al. (2024) found that the best protection of the cucumber holobiont against root-knot nematodes was achieved with communities consisting of at least six bacterial strains in the rhizosphere, which indicates the importance of a functional network as opposed to the nematode-antagonistic activity of single strains. Therefore, the functionality of such microbial processes may often depend on interactions with other organisms that co-occur in the local environment and are integrated into the holobiont’s functional network.

### Limitations

In our study, we used the tomato plant as a model for a plant holobiont. Further investigations are needed to determine the extent to which other plants exploit the emergent properties of a holobiont. Some plant species may instead employ a more autonomous ecological strategy. Differences in the strength of association between plants and their microbiomes were even found at the sub-species level in crops. Barley genotypes differed significantly in their rhizobiome-mediated defense against plant-parasitic nematodes (Elhady et al. 2021a). This could be exploited in the breeding of crops to make them more responsive to beneficial soil microbiomes. The tomato holobiont was compatible with the biotic legacy of tagetes, but not oilseed rape. Further research into other crop and cover crop combinations is needed to inform crop rotation schemes.

We only challenged the tomato holobiont with one pathogen: *M. hapla*. Further studies are needed to explore holobiont resistance against other pathogens and combinations of co-occurring pathogens. A holobiont’s strength lies in its ability to harness the diversity of functional rhizobiome networks, providing flexibility in responding to multiple pathogens and adaptability in adjusting the holobiont composition in the face of frequent biotic or abiotic challenges at a given site. As our experimental setup does not reflect the full complexity of agroecosystems, any conclusions must be confirmed by evidence from field experiments.

We only included soil from a single field in our study. The diversity of soil biota as a resource for holobiont assembly is affected by soil type, soil legacy, crop rotation schemes including cover crops, and agricultural management practices. In this study, the responsiveness to plant-conditioned nematode communities in terms of resistance of the tomato holobiont was less pronounced in the conventionally managed soil than in the organically managed soil, while the resistance of the tomato holobiont was slightly better with the soil biota from the conventional plot (first experiment). The lack of replication with respect to further pairs of differently managed field plots impedes the ability to draw conclusions. However, this result is consistent with previous research on nematode community structure in organically and conventionally managed arable soils, which found high variation but no consistent differences (Neher 1999).

### Outlook

Our study provides evidence that soil nematodes and their associated microbiota (incl. phages) are important yet previously underappreciated biotic factors that influence the emergent properties of plant holobionts and can be managed by plant-specific soil legacies. Future work should investigate the mechanisms and key taxa to understand how nematode–phage–microbiome interactions in the rhizosphere of crops can be harnessed to design more resilient and sustainable agroecosystems.

## Acknowledgements

The authors thank Olivera Topalović for thought-provoking discussions on plant holobionts and nematodes.

